# Phylostratigraphic analysis of tumor and developmental transcriptomes reveals relationship between oncogenesis, phylogenesis and ontogenesis

**DOI:** 10.1101/199083

**Authors:** Joseph X. Zhou, Luis Cisneros, Theo Knijnenburg, Kalliopi Trachana, Paul Davies, Sui Huang

**Author notes:** These authors contributed equally as first author (Joseph X. Zhou, Luis Cisneros).

## Abstract

The question of the existence of cancer is inadequately answered by invoking somatic mutations or the disruptions of cellular and tissue control mechanisms. As such uniformly random events alone cannot account for the almost inevitable occurrence of an extremely complex process such as cancer. In the different epistemic realm, an ultimate explanation of cancer is that cancer is a reversion of a cell to an ancestral pre-Metazoan state, i.e. a cellular form of atavism. Several studies have suggested that genes involved in cancer have evolved at particular evolutionary time linked to the unicellular-multicellular transition. Here we used a refined phylostratigraphic analysis of evolutionary ages of the known genes/pathways associated with cancer and the genes differentially expressed between normal and cancer tissue as well as between embryonic and mature (differentiated) cells. We found that cancer-specific transcriptomes and cancer-related pathways were enriched for genes that evolved in the pre-Metazoan era and depleted of genes that evolved in the post-Metazoan era. By contrast an opposite relation was found for cell maturation: the age distribution frequency of the genes expressed in differentiated epithelial cells were enriched for post-Metazoan genes and depleted of pre-Metazoan ones. These findings support the atavism theory that cancer cells manifest the reactivation of an ancient ancestral state featuring unicellular modalities. Thus our bioinformatics analyses suggest that not only does oncogenesis recapitulate ontogenesis, and ontogenesis recapitulates phylogenesis, but also oncogenesis recapitulates phylogenesis. This more encompassing perspective may offer a natural organizing framework for genetic alterations in cancers and point to new treatment options that target the genes controlling the atavism transition.

**One Sentence Summary:** Tracing cancer gene evolutionary ages revealed that cancer reverts to a pre-existing early Metazoan state.

## INTRODUCTION

The currently prevailing molecular-mechanistic view of cancer focuses on oncogenic mutations and considers cancer as a genetic disease. Cancer cells result from the accumulation of random somatic mutations, leading to the emergence of clones that display higher growth fitness than the normal cells^1–5^. This gene-centric paradigm, often referred to as somatic mutation theory (SMT) of cancer, has been challenged by several lines of thinking that seek to reconnect cancer biology to the organicist view of biology^6–9^. They emphasize the need to take a more integrative view, advocating that the effect of tissue-fields due to collectively acting cells, as well as non-genetic regulatory processes, such as tissue architecture remodeling^10^, angiogenesis^11^, immune-response^12^ and regeneration play central roles in tumorigenesis and tumor progression.

While the gene-centric view has now absorbed these regulatory processes by incorporating them in the list of “hallmarks of cancer” and explained their activation in tumors by genetic mutations, the organicists view these processes as the manifestation of the disruptions of developmental processes or tissue homeostasis. Although genome sequencing of human cancer have identified many driver mutations^13^, transcriptome analysis has revealed a rich dynamics of gene expression coordination that defies the simple notion of a direct linear causal relationship between an oncogenic mutations and the hallmarks of cancer^8^. Instead, such analysis points to the activation of pre-existing gene expression profiles, or “programs” that are determined by the coordinated expression behavior of distinct genes. These gene expression programs are very similar to those known to govern normal developmental processes or tissue homeostasis, such as wound-healing and regeneration. Moreover, higher-resolution genome analysis, e.g. of multiple cancer cell clones, suggest that the mutation spectrum reflect neutral evolution with only spotty evidence of selective sweeps that would support the SMT^14^. Careful experimental studies, such as those which involve lineage tracking and singe-cell resolution analysis, have provided a large body of evidences in support of non-genetic, regulated cell phenotype transitions in tumor progressions that are close variants of regulated stress-responses^15–21^. Indeed, accumulating cancer genome sequencing results suggest that some cancers do not contain recurring driving mutations at all^22,23^. On the other hand, a vast majority of premalignant genetic lesions like moles and epithelial metaplasia do not progress to invasive cancer while still carrying oncogenic mutations such as *TP53* deletion and *PI3K* or *RAS* activation^24^.

Both the gene-centric SMT paradigm and the integrative view of cancer as collective regulative disorder have in common that they seek proximate explanations^25,26^ to understand the pathogenesis of cancer in terms of altered molecular or cellular substrates and the ensuing biological processes. An epistemic alternative to this proximate explanation of the cause of cancer would be “ultimate” explanations. At this epistemic level, one idea is the cancer atavism hypothesis^27,28^. According to this view, it is unlikely for such robust, rapid and apparently well-orchestrated processes such as tumorigenesis, which require coordination of thousands of genes across many cell types, to arise simply by random mutations and selection as postulated by the SMT. Instead, cancer is hypothesized to represent a variant of a pre-existing primordial cellular state that is characterized by autonomous cell proliferation and survival capabilities, such as migration, resilience to xenobiotics and sparsity of nutrients: all the functionalities that have evolved in the context of single-cell organismal adaptation to the harsh environmental conditions during billions of years of early life evolution^27,28^. Then SMT would simply add an edge in the reactivation of such preexisting primordial programs, but not create them.

The atavism hypothesis bears semblance to a view of cancer long ago proprosed by Theodore Boveri^29^ and more recently refined by Sonnenschein and Soto that the state of unrestrained proliferation of the cancer cell is the default state of a living cell^6,7,30^. Hence cancer-causing mutations and “epigenetic” modifications only disrupt the response to signals emanating from the tissue microenvironment which normally would hold cell proliferation in check to support the cell society that underlies the multiceluar life. In the same category of more profound, “ultimate” explanations is the hypothesis by Erenpreisa and collaborators suggesting the induction of stem cell-like behaviors and even of (abortive) gametogenesis in cancer cells as response to cellular stress^31,32^, which are signatures of the activation of the life cycle characteristic of proto-metazoa organisms that occupy the evolutionary stage between unicellularity and multicellular organisms, such as hydra^33,34^ or volvox^35,36^. In these organisms, stress can stimulate the sexual reproduction cycle by activating stemness and gametogenesis to produce free living haploid unicellular cells. Furthermore, Davila and Zamorano proposed that cancer originates in cells that revert to early-stage Eukaryotic evolution, a transition caused by the deregulation of mitochondria resulting from oxidative damage to mitochondrial and nuclear DNA^37^.

These hypotheses towards “ultimate” explanantions of cancer have yet to be corroborated by solid molecular biology data but there has been a recent surge of experiental and data-driven studies exploring the relation of cancer to phylogenegic (not somatic) evolution. Chen and coworkers performed xenograft experiments of human breast cancer in immuno-deficient mice for eight generations and measured the transcriptome and genome (exome) in each generation^38^. They observed that a metastatic phenotype arose from a mutator clone representing reverse evolution driven by down-regulation of genes involved in the maintenance of metazoan multicellularity. Piermarocchi associated the centrality and the evolutionary age of each gene in the gene regulatory network derived from leukemia cells and their normal counterparts^39^. They found that slowly evolving old genes tend to interact with others of the same type, while the rapidly evolving young genes do the same with their counterparts. Domazet-Losś et al^40,41^ first introduced phylostratigraphic methods to study the evolutionary origin of cancer-associated genes and protein domains, and found that the so-called group of “caretaker genes” (involved with genome stability)^42^ were overrepresented with genes that appear among the first pre-metazoan genes, whereas the “gatekeeper genes” (controlling cellular signaling and growth processes) were associated with genes that evolved with metazoan. Cisneros et al^43^ studied gene mutation frequency as the function of evolutionary age from the ICGC whole genome sequence database^44^ and found that cancer genes were not located in the mutational ‘hot-spot’ regions of the genome, although they were more likely to be mutated than other genes. Conserved regions appeared to be functionally associated with ancient stress-induced mutagenesis programs from bacteria^43^. More recently, Trigos et al^45^ used TCGA data to analyze gene evolutionary age and transcriptome changes in cancer samples. They found that genes associated with unicellular life were up-regulated while metazoan genes were down-regulated. In addition, modules of co-expression between unicellular and multicellular genes in the GRNs were also disrupted in cancer, suggesting that human GRNs operating at the interface between multicellular and unicellular functions may be involved in the activation of primitive transcriptional programs that drive cancer. In a similar vein, Wu’s group systematically studied the karyotypes of more than 600 cancer cell lines and found that many of them jettisoned either Y or the inactive X chromosomes and that the active X often doubled with an addition of one haploid complement of autosomes – leading to a hypothesis that free-living cancer cells reconfigure their chromosomes back to a unicellular state^46^.

In spite of this progress, there are still gaps in our knowledge about the origin of cancer in terms of its evolutionary root. How do a variety of mutations and pathway activations trigger the reversion of a cell from the “contemporary” physiological state to one of the deeply-embedded hidden state? The quest for the “ultimate” explanation of the existence of cancer in terms of its evolutionary origin must also revisit the link between cancer and normal development, given the known connection between phylogenesis and ontogenesis, and the idea that cancer is a developmental disease, manifesting a maturation arrest or mis-differentiation^47,48^, as recently shown by bioinformatics analysis from Kohane el. at.^49^

In this study we obtained the evolution age of 19,177 human genes from the EGGNog phylostratigraphic database and gathered hundreds of RNA-Seq transcriptomes for ten cancer types in the TCGA database as well as their normal counterparts. After identifying genes differentially expressed in normal and cancerous tissue and comparison with their relevant gene age, we found that cancer cells typically suppress post-Metazoan genes and overexpress pre-Metazoan genes –consistent with a recently published study^45^. Compatible results were obtained by analyzing the enriched gene evolutionary ages of the relevant pathways obtained from the KEGG database. Using transcriptome data, we further determined the genes differentially expressed between human embryonic stem cells and differentiated normal epithelial cells. The result confirmed that the cell differentiation process exhibits a relationship to evolution that is opposite to that for tumorigenesis: post-Metazoan genes were overexpressed while pre-Metazoan genes were suppressed -- suggesting that malignant tumor tissue display an apparent rejuvenation, or reversal of the normal development process. Thus, if ontogenesis recapitulates phylogenesis, as biologists generally think, and if oncogenesis recapitulatesontogenesis, then our bioinformatics results now would also be consistent with the more recent idea that oncogenesis recapitulates phylogenesis, thus completing the set of bioinformatics evidence to support this triangular transitive relationship.

## RESULTS

### Oncogenes and tumor suppressor gene age distribution

Multicellular stability is the outcome of two antagonistic processes: one promotes individual cell fitness and the other promotes tissue (collective) fitness. The activities of oncogenes and tumor suppressor genes are central to the equilibrium of these two processes. Overt proliferation and cell-autonomy can result from over-activity of oncogenes that stimulate the cell division cycle, or from the reduced activity of the tumor suppressor genes. Both scenarios can be interpreted as biological functions that support single cellular fitness and can result from activating mutations in the former or inactivating mutations in the latter.

Accordingly, what we now call tumor suppressor genes have been proposed to have played a critical role in the evolution of multicellularity^50–54^, as part of the complex machinery of top-down controls necessary for establishing organismal coherence and homeostatic stability. On the other hand, retaining oncogenes offers a less apparent general advantage to multicellular organisms. Given that cell proliferation could be a default state in unicellular organism^55^, as long as there is sufficient energy supply and space, one possible reason to retain oncogenes is that extant oncogenes evolved from genes involved in the machinery that implements this default state, or are derived from ancestral ‘contingency genes’ of unicellular organism that originally evolved to enable populations of cells to respond to stressful environments by promoting proliferation. In multicellular organism then such genes would have evolved to promote tissue expansion during development (providing the somatic cells) and regeneration (repairing the somatic cells) – both would serve the propagation of the germline. In any case, one can expect that evolution of a fraction of oncogene precursor genes predate the appearance of multicellularity or at least, that of tumor suppressor genes.

We first used existing knowledge bases to define three set of genes: all human genes, genes that have been labeled ‘tumor suppressor genes’ and genes that have been labeled ‘oncogenes’^56,57^. In parallel, we categorized all genes in five evolutionary age groups based on the eggNOG database^58^, as shown in Fig.1a, from old to young: LUCA (last unicellular common ancestor), Eukaryota, Metazoa, Vertebrata and Primata (for details about the determination of gene ages see Material and Method). When multicellular life appeared on earth, the cellular traits that are heir to unicellular protozoa were suppressed or tightly controlled by regulatory mechanisms which evolved to promote collective over individual fitness^59^. Since evolution tends to build layers of phenotype on top of existing ones by amending developmental processes and suppressing primitive elements^47,60^, a concept that explains the existence of atavism (such as tails in human)^61^, cancer could be depicted as a form of cellular atavism: a reversion to single cell behavior due to reactivation of hidden (normally suppressed) ancient “genetic programs” (Fig. 1b). This reactivation could be triggered by mutations (or other disturbances) that disrupt the functional mechanisms that protect multicellularity^41,43,45^. Recent work^43^ showed that ancient mutational responses to stress, co-opted in multicellular life to maintain diversity in the germline and immune system, can cause cells to revert to a primordial form that displays the cancer phenotype, including elevated genetic instability and accelerated somatic adaptation.

**Figure 1.**
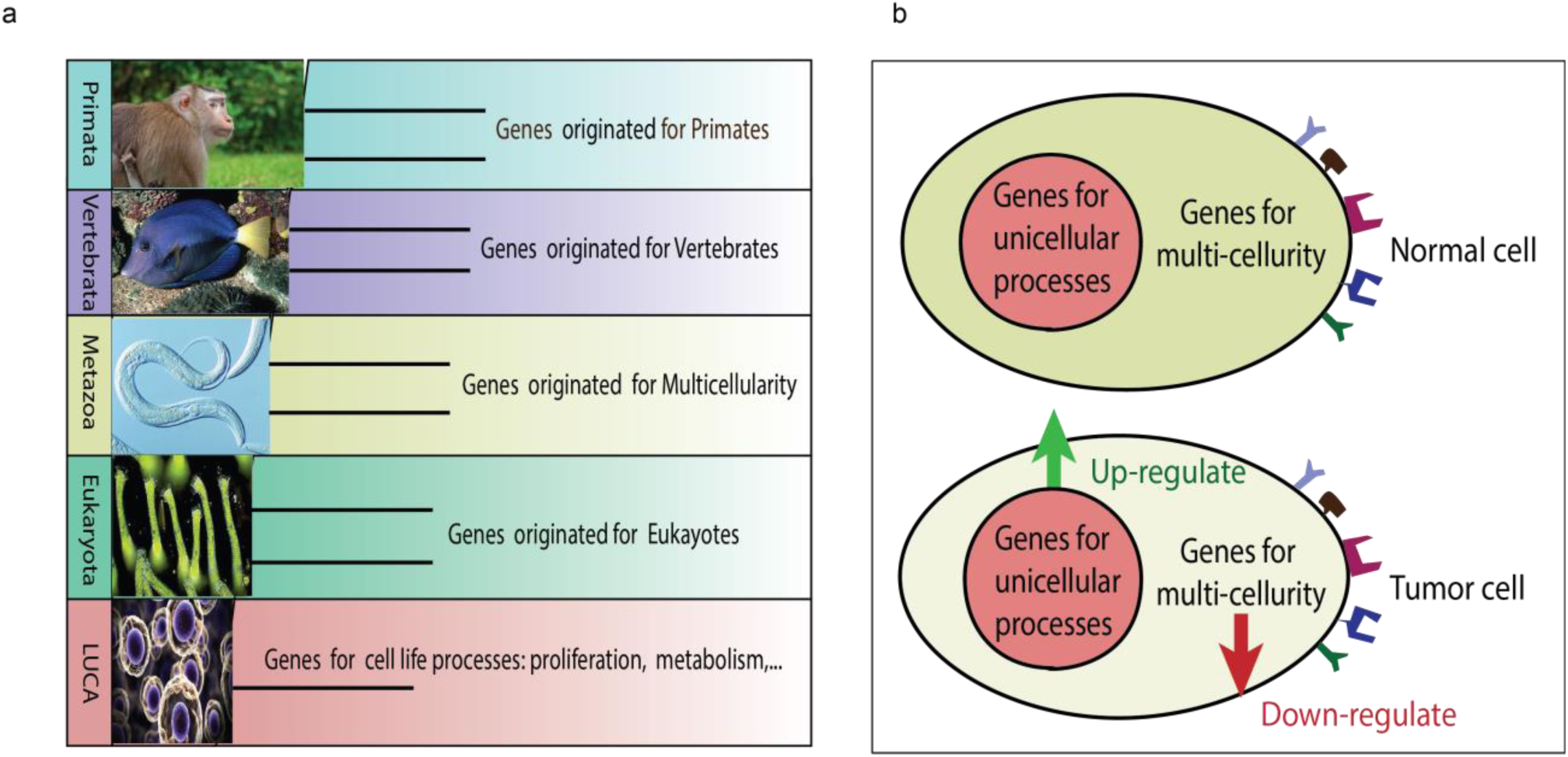
The scheme of gene evolution and the cancer atavism hypothesis. (**a**) Gene orthologues were extracted from the eggNOG database. We identify all orthologous groups containing human genes and reduced taxonomic levels to five basic levels: LUCA, Eukaryota, Metazoa, Vertebrata and Primata. (**b**) Cancer atavism hypothesis: cancer is a pre-programmed state, which enable cell growth and highly efficient adaptation to environmental changes, honed by a long period of evolution in ancestral life and subsequently suppressed in multicellular life. In the cancer phenotype, genes that play a role in single-cellular processes are up-regulated while genes that play a role in multicellular process are down-regulated compared with the normal tissue.

In a first analysis, we counted the number of genes with the labels ‘tumor suppressor’ or ‘oncogene’ and computed their relative frequency among the gene classes of the different evolutionary age and compared it with the frequency of all genes as the background, i.e., we determined the relative enrichment and depletion of suppressor and oncogenes in each of the evolutionary age groups. Fig. 2a presents a simple comparison of the frequencies of occurrence in each evolutionary age group for the three sets of genes, all human genes, tumor suppressor genes and oncogenes. In Fig. 2b we use an enrichment score based on an adjusted residual analysis (see Material and Methods), comparing suppressor genes and oncogenes against all human genes as the background. Both plots show that the frequency of the tumor suppressor genes are depleted for genes at the Vertebrata level while they are enriched in the age groups of the Eukaryota and Metazoa level. Oncogenes, however, also display a stronger enrichment for genes of the Eukaryota and Metazoa levels. By contrast, the frequency of oncogenes are specifically depleted for genes in Vertebrata and Primata levels. Thus in general both gene sets follow a very similar pattern: both groups of genes mutated in cancer typically evolved before or during the evolution of multicellularity, but oncogenes seem to have a stronger pre-metazoan component earlier, appearing during the evolution of eukaryotes, as suggested by the fact that their frequency is actually depleted in LUCA.

**Figure 2.**
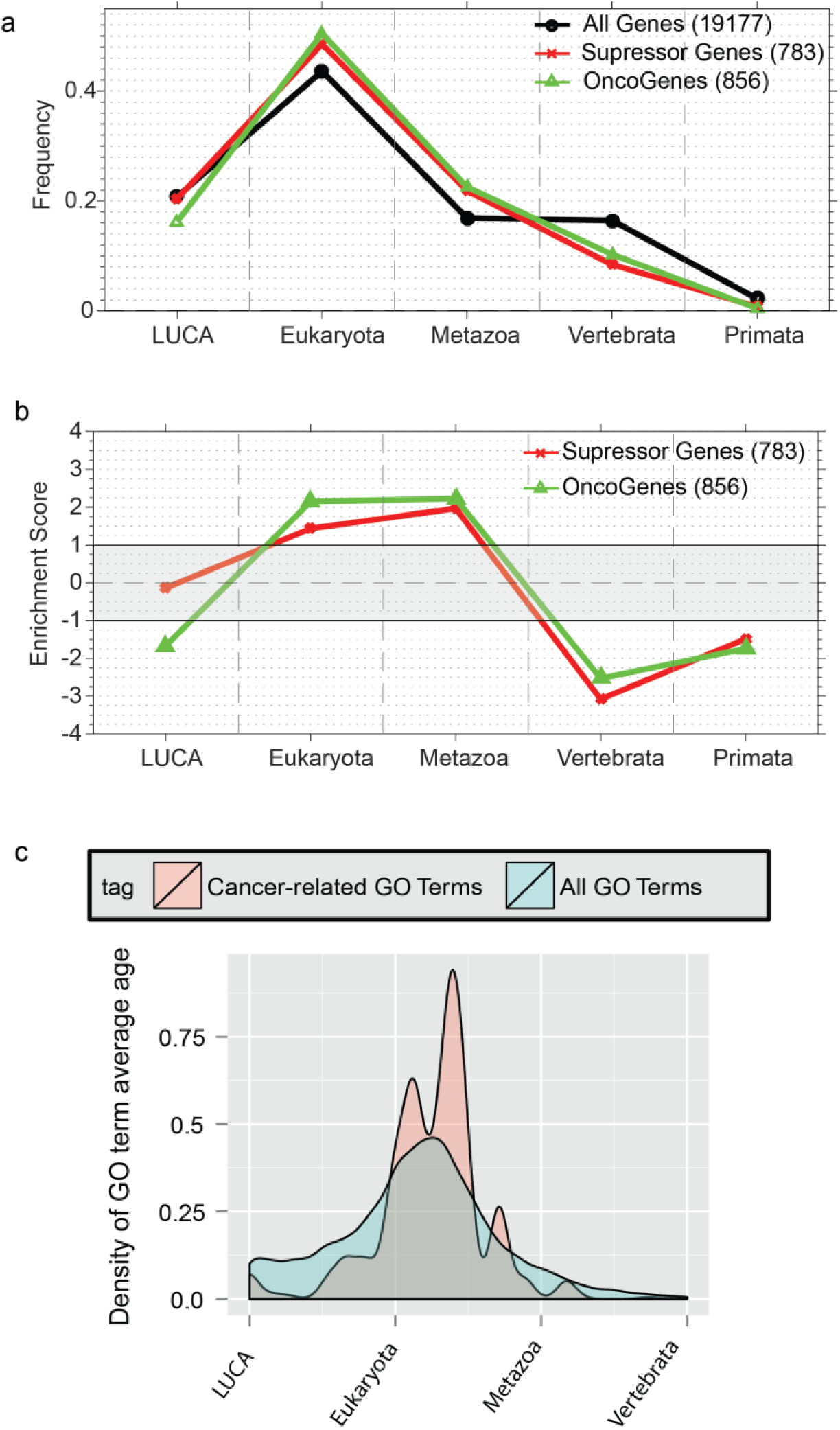
The age distribution of the annotated cancer genes and cancer GO terms. (**a**) The age distribution for the whole set of 19,177 human genes (black circles) with phylostratigraphic age and two relevant gene sets in cancer genomics: tumor suppressor genes (red exes) and oncogenes (green triangles). (**b**) We present the age enrichment score for these two gene sets using all genes as the background. The values that fall outside of the grey band are statistically significant with p-value < 0.05. It shows that both tumor suppressor genes and oncogenes are over-represented in Eukayota and Metazoa and under-represented in Vertebrata. Additionally, Oncogenes are under-represented in LUCA. (**c**) The age distribution of the cancer-related GO terms compared with all GO terms (data accessed on 07/2017 from Gene Ontology Consortium). The average ages of all cancer-related GO terms are compared with those of all GO terms, which shows that the cancer-related GO terms are enriched between Eukayota and Metazoa. This GO term age analysis agrees with the result of cancer gene age analysis.

Since genes function as interacting gene networks, we also examined the gene sets involved in cancer based on gene ontology (GO) annotations which places genes with the same function into groups based on shared GO labels. In Fig. 2c we separately curated all GO terms and cancer-related GO terms and then calculated the average age of each one using our gene age database. The cancer-related GO terms display a strong enrichment in the age groups between Eukaryota and Metazoa, again suggesting that cancer-related genes evolved before or during the evolution of multicellularity.

### The gene age distribution of cancer-related and house-keeping pathways

Since there is no significant difference in the evolutionary age enrichment between oncogenes and tumor suppressors found in tumor genomes, it has been suggested that considering pathways as the “units” targeted by mutations may be more appropriate^62^. Therefore, we next curated cancer-related pathways from the KEGG database^63^ and analyzed the evolutionary gene age distribution enrichment by comparing the age distribution of the gene sets associated with a given pathway with the background distribution of non-cancer genes. We analyzed 25 pathways associated with cancer (see Table 1) as well as 14 housekeeping pathways (see Table 2) for their enrichment in the evolutionary gene age groups. The latter set can be used to establish a benchmark for the comparison of the patterns observed in cancer.

**Table 1.** Cancer related Pathways from KEGG.

| Name | KEGG Gene Numbers | KO Class | KO Subclass | BRITE category label |
| --- | --- | --- | --- | --- |
| Cell cycle | 124 | Cellular Processes (hsa09140) | Cell growth and death (hsa09143) | hsa04110 |
| p53 signaling pathway | 68 | Cellular Processes (hsa09140) | Cell growth and death (hsa09143) | hsa04115 |
| Apoptosis | 138 | Cellular Processes (hsa09140) | Cell growth and death (hsa09143) | hsa04210 |
| Focal adhesion | 199 | Cellular Processes (hsa09140) | Cellular community - eukaryotes (hsa09144) | hsa04510 |
| Cell adhesion molecules (CAMs) | 145 | Cellular Processes (hsa09140) | Cellular community - eukaryotes (hsa09144) | hsa04514 |
| Adherens junction | 72 | Cellular Processes (hsa09140) | Cellular community - eukaryotes (hsa09144) | hsa04520 |
| MAPK signaling pathway | 255 | Environmental Information Processing (hsa09130) | Signal transduction (hsa09132) | hsa04010 |
| ErbB signaling pathway | 86 | Environmental Information Processing (hsa09130) | Signal transduction (hsa09132) | hsa04012 |
| mTOR signaling pathway | 151 | Environmental Information Processing (hsa09130) | Signal transduction (hsa09132) | hsa04150 |
| PI3K-Akt signaling pathway | 342 | Environmental Information Processing (hsa09130) | Signal transduction (hsa09132) | hsa04151 |
| TGF-beta signaling pathway | 84 | Environmental Information Processing (hsa09130) | Signal transduction (hsa09132) | hsa04350 |
| RAS Signaling Pathway | 227 | Environmental Information Processing (hsa09130) | Signal transduction (hsa09132) | hsa04014 |
| HIF-1 signaling pathway | 100 | Environmental Information Processing (hsa09130) | Signal transduction (hsa09132) | hsa04066 |
| WNT signaling pathway | 143 | Environmental Information Processing (hsa09130) | Signal transduction (hsa09132) | hsa04310 |
| Notch signaling pathway | 48 | Environmental Information Processing (hsa09130) | Signal transduction (hsa09132) | hsa04330 |
| Hedgehog signaling pathway | 47 | Environmental Information Processing (hsa09130) | Signal transduction (hsa09132) | hsa04340 |
| VEGF signaling pathway | 59 | Environmental Information Processing (hsa09130) | Signal transduction (hsa09132) | hsa04370 |
| Jak-STAT signaling pathway | 156 | Environmental Information Processing (hsa09130) | Signal transduction (hsa09132) | hsa04630 |
| Cytokine-cytokine receptor interaction (CytK-CytK) | 270 | Environmental Information Processing (hsa09130) | Signaling molecules and interaction (hsa09133) | hsa04060 |
| ECM-receptor interaction | 82 | Environmental Information Processing (hsa09130) | Signaling molecules and interaction (hsa09133) | hsa04512 |
| Mismatch Repair (MMR) | 23 | Genetic Information Processing (hsa09120) | Replication and repair (hsa09124) | hsa03430 |
| Homologous Recombination (HR) | 41 | Genetic Information Processing (hsa09120) | Replication and repair (hsa09124) | hsa03440 |
| PPAR signaling pathway | 72 | Organismal Systems (hsa09150) | Endocrine system (hsa09152) | hsa03320 |
| Melanogenesis (Melano) | 101 | Organismal Systems (hsa09150) | Endocrine system (hsa09152) | hsa04916 |
| Transcriptional misregulation in cancer (TMR) | 186 | Human Diseases (hsa09160) | Cancers (hsa09161) | hsa05202 |

Source: KEGG Pathways Database: 05200 - Pathways in cancer.
http://www.genome.jp/kegg-bin/show_pathway?org_name=hsa&mapno=05200

**Table 2.**
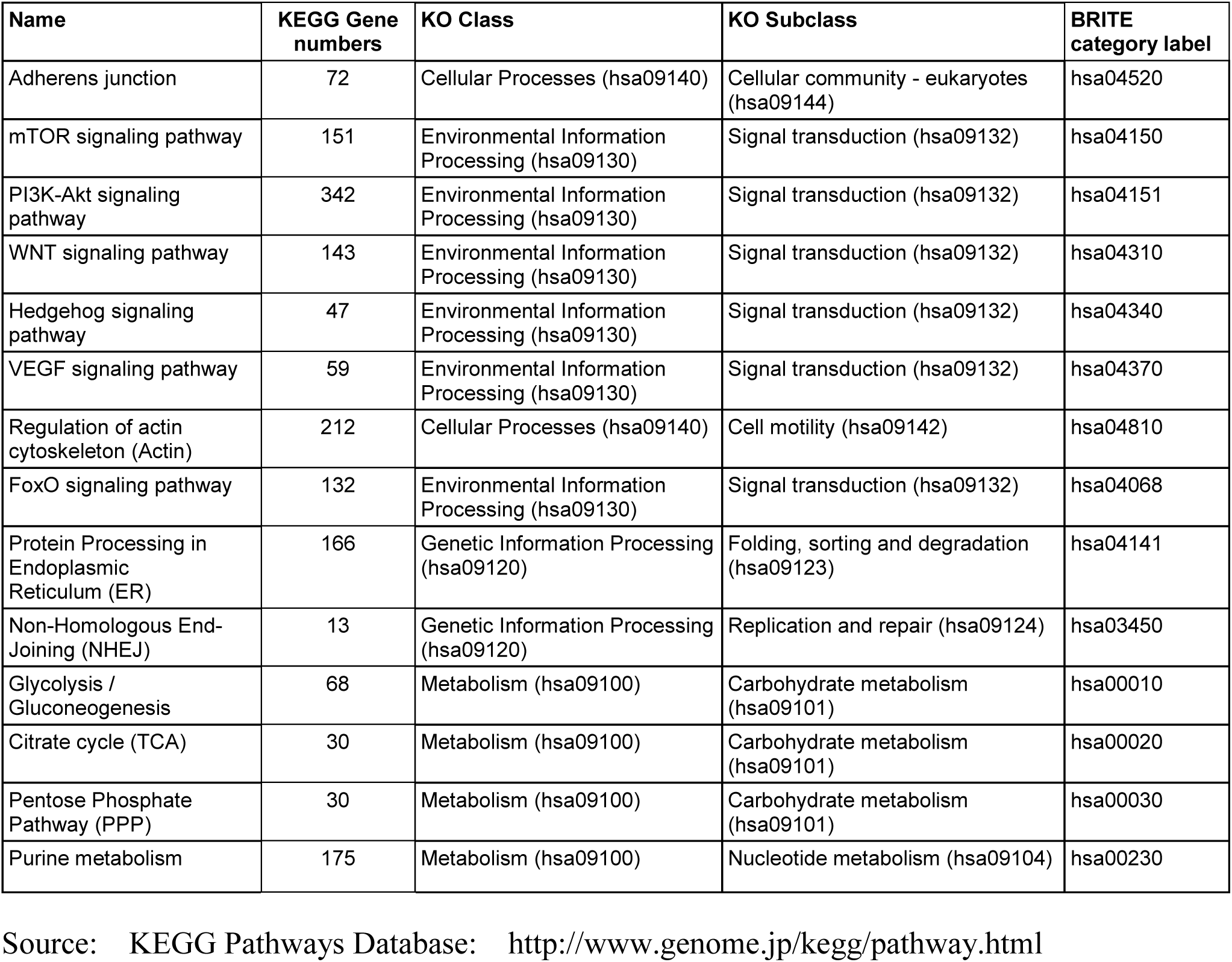
House-keeping pathways from KEGG.

| Name | KEGG Gene numbers | KO Class | KO Subclass | BRITE category label |
| --- | --- | --- | --- | --- |
| Adherens junction | 72 | Cellular Processes (hsa09140) | Cellular community - eukaryotes (hsa09144) | hsa04520 |
| mTOR signaling pathway | 151 | Environmental Information Processing (hsa09130) | Signal transduction (hsa09132) | hsa04150 |
| PI3K-Akt signaling pathway | 342 | Environmental Information Processing (hsa09130) | Signal transduction (hsa09132) | hsa04151 |
| WNT signaling pathway | 143 | Environmental Information Processing (hsa09130) | Signal transduction (hsa09132) | hsa04310 |
| Hedgehog signaling pathway | 47 | Environmental Information Processing (hsa09130) | Signal transduction (hsa09132) | hsa04340 |
| VEGF signaling pathway | 59 | Environmental Information Processing (hsa09130) | Signal transduction (hsa09132) | hsa04370 |
| Regulation of actin cytoskeleton (Actin) | 212 | Cellular Processes (hsa09140) | Cell motility (hsa09142) | hsa04810 |
| FoxO signaling pathway | 132 | Environmental Information Processing (hsa09130) | Signal transduction (hsa09132) | hsa04068 |
| Protein Processing in Endoplasmic Reticulum (ER) | 166 | Genetic Information Processing (hsa09120) | Folding, sorting and degradation (hsa09123) | hsa04141 |
| Non-Homologous End-Joining (NHEJ) | 13 | Genetic Information Processing (hsa09120) | Replication and repair (hsa09124) | hsa03450 |
| Glycolysis / Gluconeogenesis | 68 | Metabolism (hsa09100) | Carbohydrate metabolism (hsa09101) | hsa00010 |
| Citrate cycle (TCA) | 30 | Metabolism (hsa09100) | Carbohydrate metabolism (hsa09101) | hsa00020 |
| Pentose Phosphate Pathway (PPP) | 30 | Metabolism (hsa09100) | Carbohydrate metabolism (hsa09101) | hsa00030 |
| Purine metabolism | 175 | Metabolism (hsa09100) | Nucleotide metabolism (hsa09104) | hsa00230 |

Source: KEGG Pathways Database: http://www.genome.jp/kegg/pathway.html

In each case, cancer pathways were grouped according to their KEGG-BRITE orthologous groups, which spans categories ranging from those responsible for essential cell functions, such as metabolism, genetic information processing and cellular processes, to those of organismal-level processes, such as environmental information processing, organismal systems and a few categories involved in human diseases.

Among the cancer pathways (Fig. 3a), our analysis shows that pathways associated with cellular growth and genetic information processing, which could be considered cell-level processes, are typically enriched in the ancient LUCA and Eukaryota gene groups, while most pathways controlling the organismal-level processes are enriched in the group of genes that appeared with Metazoa and Vertebrata. House-keeping pathways (Fig. 3b) related to metabolism are enriched in the group of LUCA genes while the pathways involved in signal transduction again, are enriched in the groups of Metazoa and Vertebrata genes. Thus, the functional distinction between cell-level and organismal-level functions is consistent with the gene age profile.

**Figure 3.**
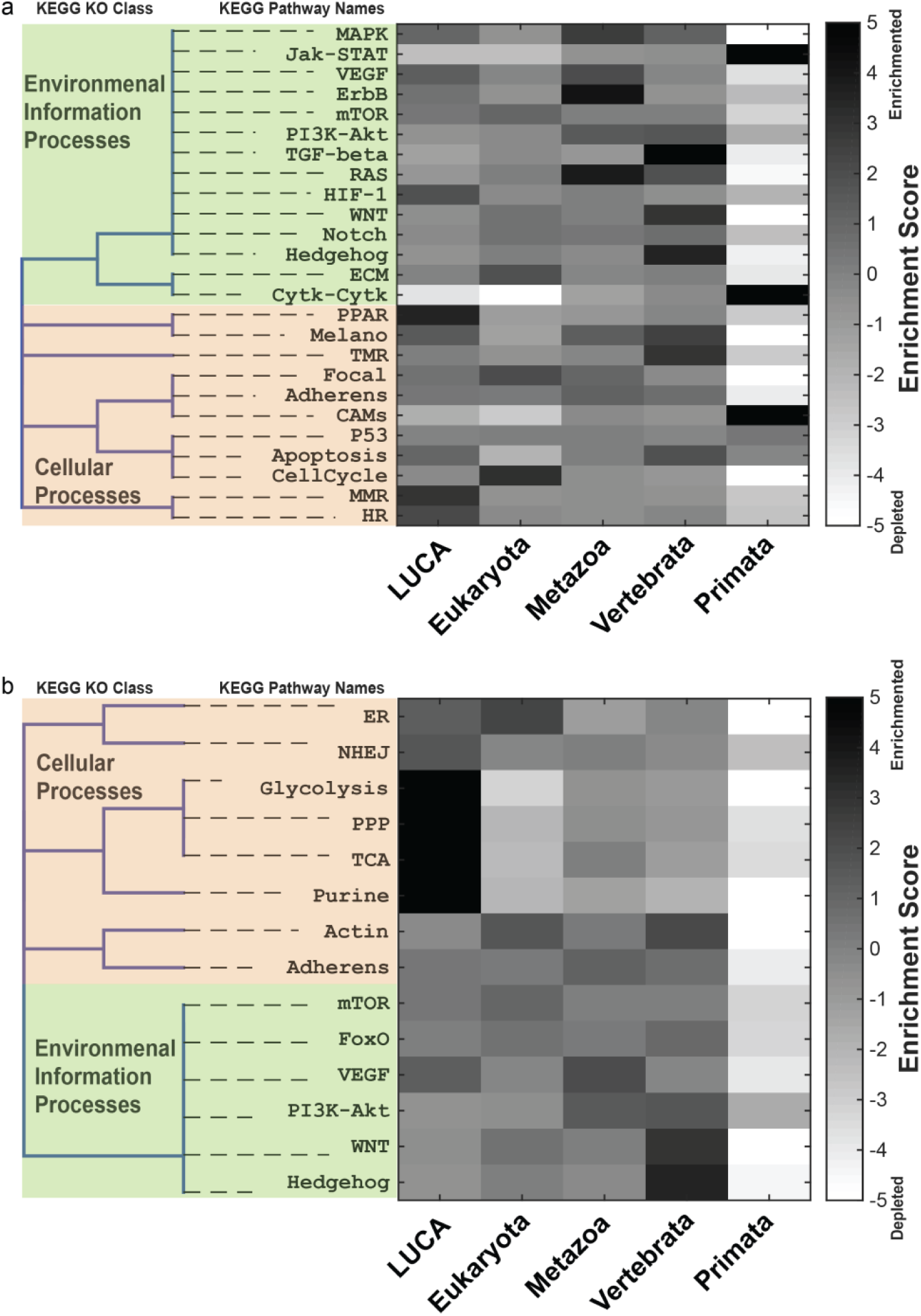
Age enrichment of cancer-related pathways and house-keeping pathways. Pathways from KEGG database are grouped according to their KEGG BRITE Orthologous classes (**a**) In cancer-related pathways (see Table 1), the categories associated with cellular growth and genetic information processing (such as Cell cycle, *mTOR, HIF1a, PPRA, MMR* and *HR*) are typically enriched in old ages (LUCA and Eukaryota), while most pathways controlling the multicellular processes, like signal transduction pathways (*ErbB, TFGbeta, RAS, WNT, Notch* and Hedgehog etc.), are enriched in Metazoa and Vertebrata evolutionary levels; (**b**) For house-keeping pathways (see Table 2), the categories related to metabolism (such as Glycolysis, PPP, TCA, mToR and Purine) are enriched in LUCA genes while the ones involved in signal transduction (Actin, Adherens junction, *VEGF, PI3K, WNT* and *Hedgehog* etc.) are enriched in Metazoa and Vertebrata

It is important to mention that enrichment in this analysis only pertains to the number of genes (see details in Table 1) in a pathway that falls in each age category, and thus this is a crude analysis: pathways constitute of genes connected by a regulatory relationship and as such, different subsets of genes do not necessarily have the same relevance in the dynamics and functionality of each pathway. Therefore, the enrichments that we observe are just a testament of the overall trend in the fraction of genes that evolved before or after multicellularity. The changes in the activities and topologies of these pathways could of course be equally important to a full understanding of the evolution of their biological functions, but we still obtain fairly consistent results from the gene membership annotation alone.

### Differentially expressed genes between normal and tumor tissues show divergent enrichment of gene evolution age

Beyond looking at the evolutionary age profile of genes and pathways from curated knowledge bases that predetermines relationship to cancer, we next analyzed the characteristic gene expression profiles directly from cancer tissue samples as has recently been done by Trigos et al^45^. We used the currently largest cancer omics database, TCGA, which contains thousands of primary tumor RNA-Seq samples and their corresponding normal tissue samples for various cancer types.

To define a gene as differentially expressed between cancer and normal tissues in view of the vast heterogeneity of the transcriptomes even among the samples within the same tumor type (“inter-patient heterogeneity”) we first performed a standard t-test on each cancer sample against the population of corresponding normal tissues. If a gene expression value in a cancer sample is three standard deviations σ^n^ above the population mean 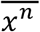 for the corresponding normal tissue, this gene is deemed “overexpressed”, i.e. if 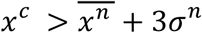, then gene χ^c^ is “overexpressed” in this cancer tissue sample; conversely, 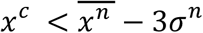, gene χ^c^ is considered “suppressed” in cancer tissue sample. Second, the overlap frequency f, as the percentage of samples that share the commonly suppressed or overexpressed gene, was used as another criterion for categorization. For instance the set of up-regulated genes with f > 10% consists of all the genes that are overexpressed in more than 10 percent of the studied samples, and so forth, with higher % values indicating more robust alteration of expression in cancer.

We then proceeded to study the evolution age distribution enrichment of overexpressed and suppressed genes with different overlap frequencies *F*.

We first performed this analysis for breast cancer samples and the corresponding normal tissue samples. As shown in Fig. 4a, with regard to the evolutionary age group, the frequency of the genes that exhibited most pronounced differential suppression in breast cancer were significantly enriched for Metazoa and Vertebrata evolutionary ages while slightly depleted in the Eukaryota age group. This result suggests that malignancies tend to suppress the expression of post-metazoan genes, those likely involved in multicellular functions such as intercellular signaling. At the same time, the more primordial genes, likely involved in essential cell functions, are typically not down-regulated in tumors. Similarly in Fig. 4b, the frequencies of genes overexpressed in cancer were significantly enriched with genes of the LUCA level age group and depleted of genes of the Vertebrata age group. Thus, the overall trend observed was that genes up-regulated in breast cancer are typically ancient, although not consistently up-regulated across all samples.

**Figure 4.**
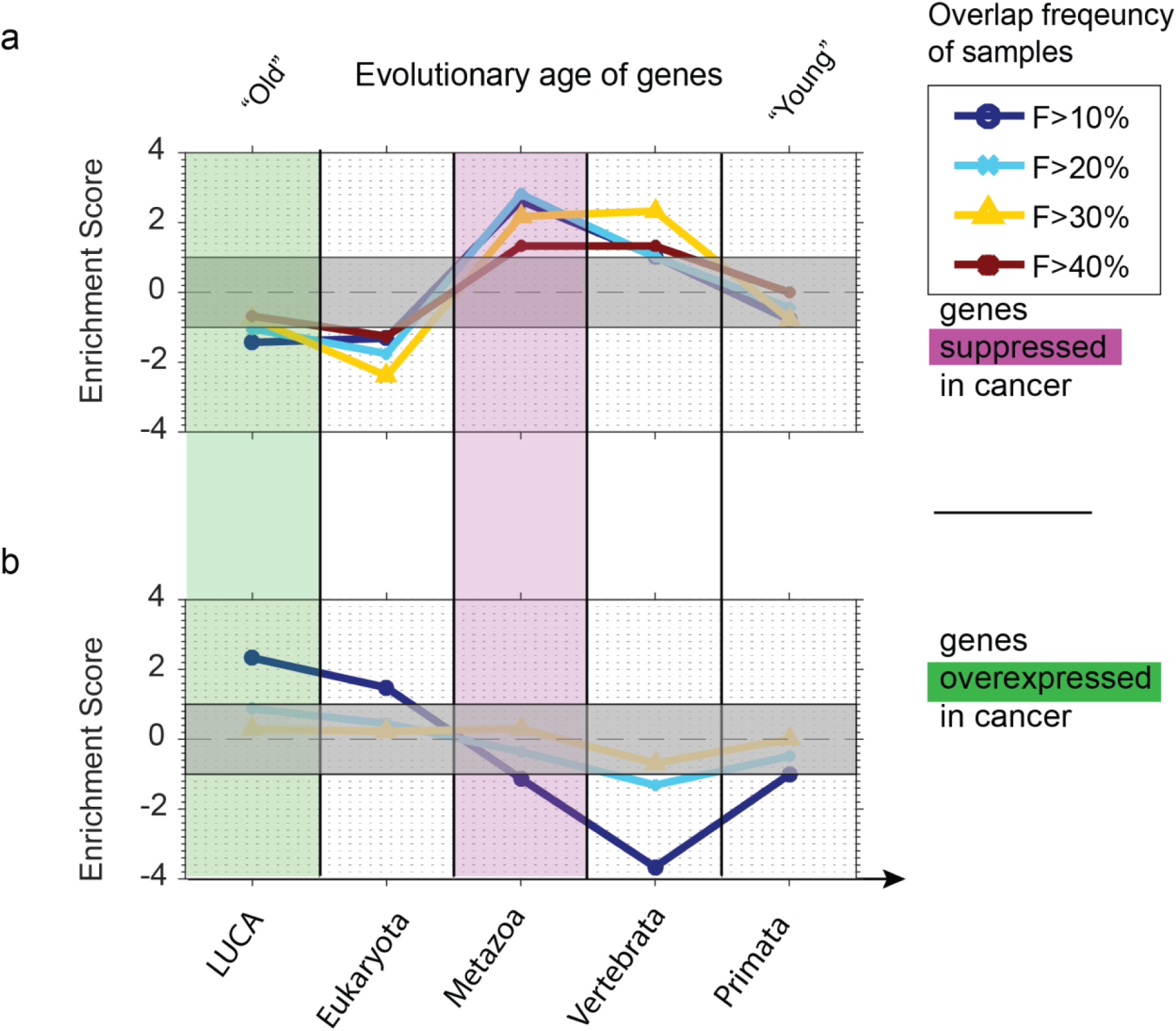
Evolutionary age enrichment analysis of genes suppressed and overexpressed in breast cancer compared with normal breast tissue from TCGA transcriptome data. (**a**) The frequency of genes suppressed in breast cancer is significantly enriched in Metazoa and Vertebrata evolutionary ages and is evidently depleted of pre-metazoan evolutionary age for various *F* values (indicating the % of samples that share the overexpression/suppression in cancer). This trend seems to be most significant for genes whose suppression in cancer was defined at *F*>30%, and is less significant at *F*>40% because the number of differentially expressed genes becomes too small. (**b**) Overexpressed genes do not seem to have a strong enrichment in any evolutionary age except for the set of genes with *F*>10%; these genes were significantly enriched with LUCA evolutionary age and depleted of post-metazoan evolutionary age (No genes were up-regulated with F>40%).

A similar analysis was performed for nine other cancer types (see details about cancer types in Material and Method Section) for genes with a sample overlap frequency taken at F>30% in all cases in order to focus on genes with consistent overexpression or suppression patterns across the cancer data (F>30% gave most consistent results in the case of breast cancer). The frequencies of genes of a given evolutionary age that were suppressed in cancer were typically higher for the post-metazoan genes and reduced for pre-metazoan genes in almost all cancer types studied (see Fig. 5a). Notable exceptions were Glioblastoma (GBM) and Kidney Renal Clear Cell Carcinoma (KIRC) which were significantly depleted of the Vertebrata age group, and Head-Neck Squamous Cell Carcinoma (HNSC) and Uterine Corpus Endometrial Carcinoma (UCEC) which displayed no significant evolutionary age enrichment. Interestingly, genes suppressed in Glioblastoma (GBM) were enriched in the Eukaryota age group. This reversal compared to other cancer types seems to be driven by an enrichment of genes functionally involved in cell adhesion, calcium ion binding and cadherin specific to the tissue type. Conversely, as shown in Fig. 5b, the genes overexpressed present a more regular behavior: there was an enrichment in the LUCA age group and depletion in the post-metazoan age groups for most cancer types. Kidney Renal Clear Cell Carcinoma (KIRC) again was an outlier in that overexpressed genes displayed enrichment in the Vertebrata age group. This set of genes was enriched with genes associated with membrane functions and production of immunoglobulins, also a specific property of the tissue type

**Figure 5.**
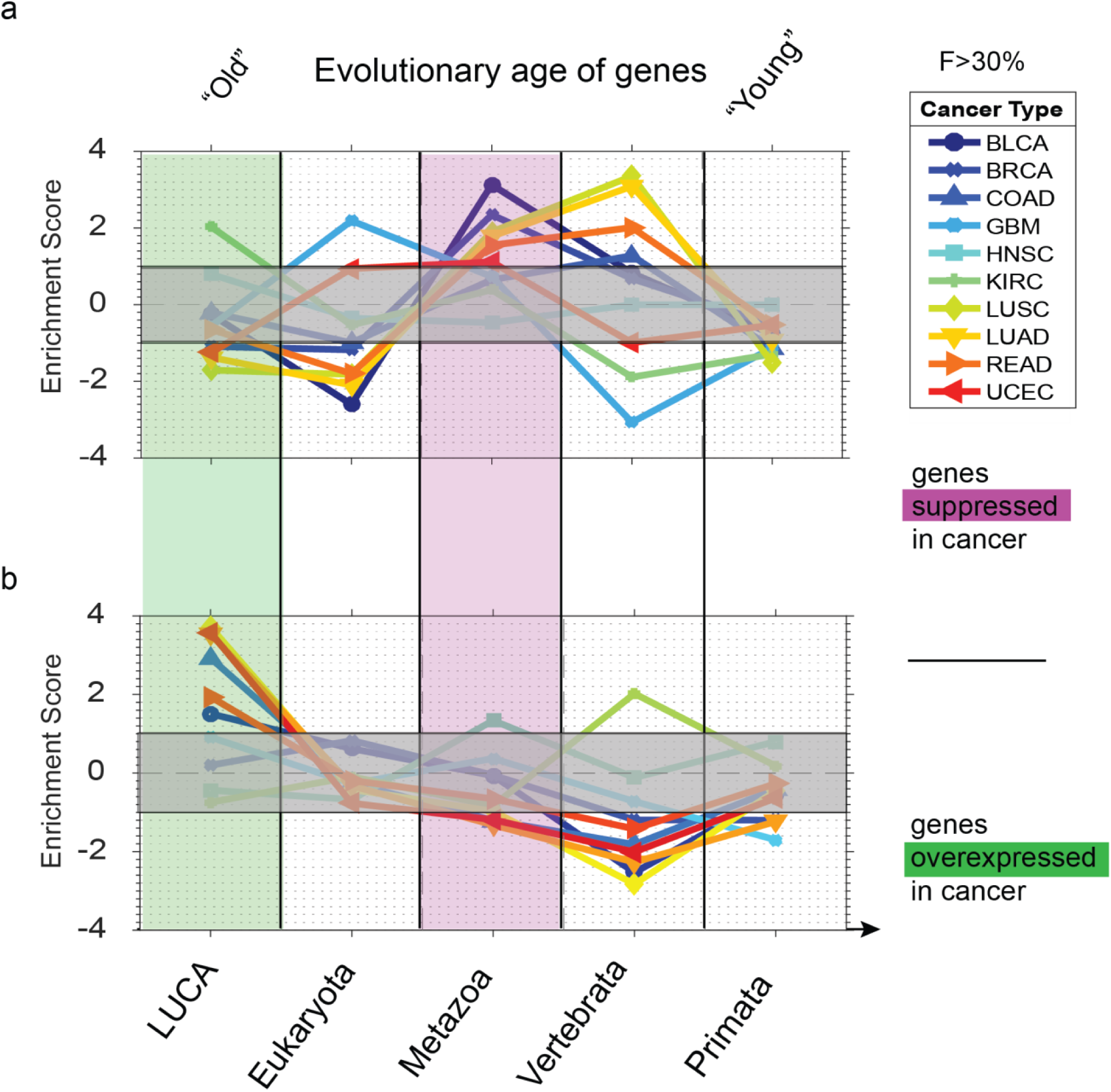
Evolutionary age enrichment analysis of genes suppressed and overexpressed in ten cancer types from TCGA transcriptome data. (**a**) Suppressed genes are typically enriched with post-metazoan genes and depleted of pre-metazoan genes in almost all studied cancer types (see Table 3). (**b**) The overexpressed genes present a more consistent behavior: there is enrichment in LUCA ages and depletion of post-metazoan ages for most cancer types. Kidney Renal Clear Cell Carcinoma (KIRC) is again an outlier presenting enrichment in Vertebrata ages.

**Table 3.**
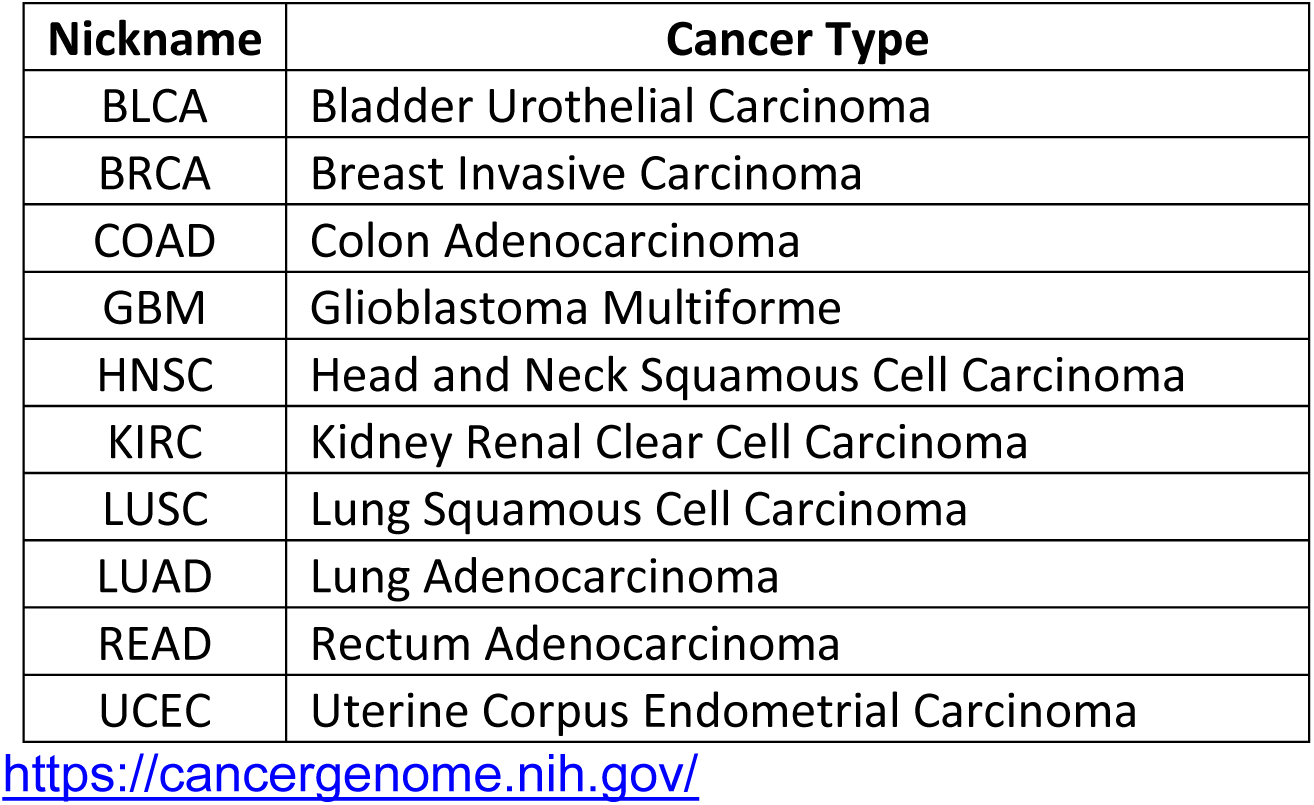
Cancer types from TCGA.

In conclusion, the enrichment of cancer-associated genes with respect to their evolutionary age in ten cancer types from the TCGA RNA-Seq transcriptome database suggests that cancer cells tend to suppress the expression of post-metazoan genes and overexpress pre-metazoan genes. These results are in line with the analysis of oncogenes vs. suppressor genes and suggest that cancer cells acquire a state that is close to a primordial state defined by expression of genes that existed in pre-metazoan stage of evolution, as predicted by the cancer atavism hypothesis.

### Biological processes involved in differentially expressed genes between normal and tumor tissues

Besides knowing the most frequent evolutionary age of genes that are overexpressed and suppressed in cancer tissues, it is also desirable to identify their biological functions of cancer-associated genes for which we had analyzed the evolutionary age. We computed the enrichment of GO terms for the differentially expressed genes with overlap frequency *F* > 30% for each cancer type. From Fig. 6a it is evident that overexpressed genes in cancer are involved with basic cell processes such as cell division, cell cycle (cell cycle phase, mitotic cell cycle etc.) and organelle fission (including spindle, cytoskeleton and nuclear division). On the other hand, the suppressed genes in cancer are related to multicellular development, embryonic organ development, morphogenesis, cell differentiation and inter-cellular signal transmission, as shown in Fig. 6b.

**Figure 6.**
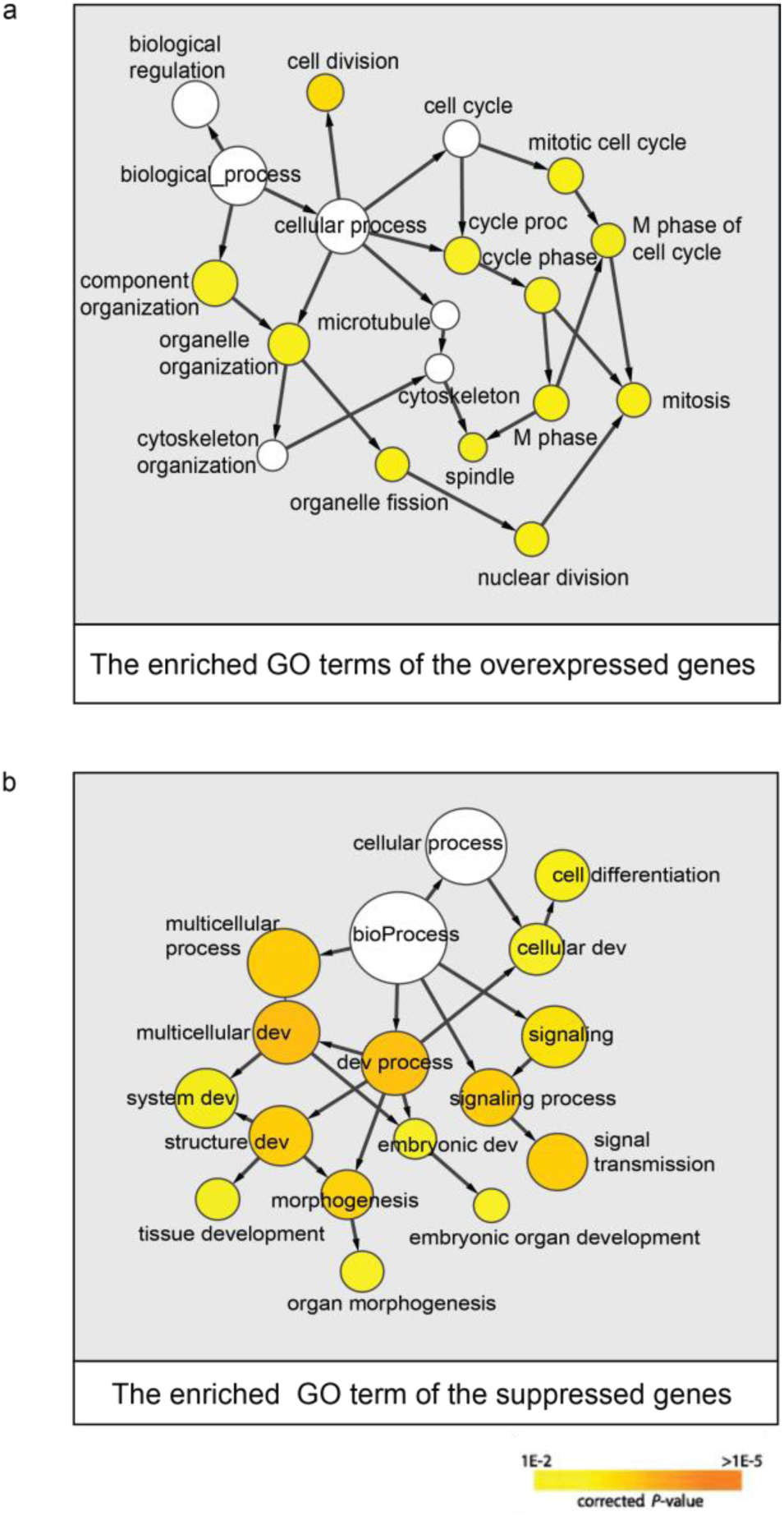
Enriched GO terms of the genes commonly overexpressed and suppressed cancer sample transcriptomes compared with that of normal tissues. We associated the biological functions to the genes commonly overexpressed and suppressed in the tumor samples across 10 cancer types compared with their normal tissues. The circle size represents the gene number within a GO term. The arrows represent the hierarchical relationships between GO terms. (**a**) Overexpressed genes in cancer samples are enriched in GO terms such as cell division, cell organelles fission (spindle, cytoskeleton, nuclear etc.), cell cycle process (mitosis, M phage) etc.; (**b**) Suppressed genes in cancer samples are enriched in GO terms such as embryonic organ development process, tissue development, morphogenesis, multicellular process, cell differentiation and inter-cellular signal transmission.

This functional analysis, which agrees with a vast body of work on GO enrichment among cancer genes in the past^64–70^, is in line with our results of the evolution age enrichment analysis: cancer cells overexpress old genes mainly involved with essential cell functions, such as cell cycle and cell division, and suppresses young genes typically associated with multicellular functions, such as cell differentiation and inter-cellular signaling. This agrees with the observation that an early characteristic of cancer cells the loss the differentiated cell identify and reversion to a stem-cell-like state^71^. Specific mutational patterns of oncogenes and tumor suppressors are not necessary conditions for cells to achieve a cancer cell state, explaining why patients with the same disease can have totally different spectra of gene mutations and overlap minimally with each other. Since the gene regulatory network is a complex system, there are multiple ways to reach the cancer cell state from many distinct combinations of gene mutations or disrupted regulation controls. Therefore, genes that are mutated in cancer display a large diversity while the implementations of the cancerous cell states share a typical core functionality, namely, suppressing the specific functions of cell differentiation and inter-cellular signaling while enhancing the functions of cell division cycle.

### Genes differentially expressed between embryo stem cells and the differentiated breast cells show the divergent enrichment of gene evolution age

If tumorigenesis is a process in which cells revert to a well-programmed early-evolved stem-cell-like state, it is very tempting to look at the normal cell differentiation process and align the oncogenesis-phylogenesis axis of analysis performed above with that of the long recognized oncogenesis-ontogenesis axis. If a similar mechanism is at work, we should see that the differentiation process runs opposite to that of tumorigenesis: differentiated cells will suppress the evolutionarily “old” genes involved with essential cell functions such as cell cycle and cell division but overexpress “young” genes which are associated to the multicellular processes and inter-cellular signaling.

We curated microarray gene expression profiles from 123 samples of embryonic stem cells (ESC) and from 159 breast epithelial cell lines of primary cell cultures from GEO (see details in Material and Methods). After applying quantile normalization to all microarray data, we again used the standard *t-*test to select the set of genes exhibiting differential expression between breast epithelium and ESC: if a gene *X*_B_ in the breast cell transcriptome is expressed at three standard deviations σ_ESC_ above the population mean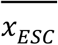 of the ESC, this gene is considered as overexpressed. Thus, if 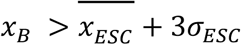, gene *x*_B_ is overexpressed. Conversely if 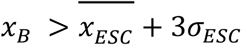 gene *x* _B_ is suppressed. Because even false discovery rate for upregulated genes is 5%, our enrichment scores are still far above our enrichment criteria (p<0.01). Thus, we don’t need to perform multiple test correction in the method of selection of overexpressed and suppressed genes. The overlap frequency of samples *F* is the percentage of samples that share the commonly suppressed / overexpressed gene.

As shown in Fig. 7a, the frequency of the genes suppressed in breast cells were significantly enriched in the LUCA and Eukaryota age groups and depleted in the Metazoa, Vertebrata and Primata age groups. By contrast, genes overexpressed in these epithelial cells were enriched in the Metazoa and Vertebrata age groups while depleted in pre-Metazoan age groups (Fig. 7b). Thus, comparing epithelial cells (mature) to embryonic cells (immature) as opposed to comparing cancer cells (presumed immature) to their normal counterpart (mature) yielded opposite results for the evolutionary age of the genes suppressed/overexpressed in these two comparison pairs. Since here the transcriptomes were mostly measured from the same or similar cells, the diversity of cells does not play as much a role as was the case for the comparison between cancer and normal tissues. This may explain why the enrichment scores vary little with respect to different overlap frequencies *F*. In conclusion, human breast cell differentiation seems to have the opposite effect on evolutionary age of the genes expressed compared to from tumorigenesis. To reach the fully differentiated state, cells tend to suppress the pre-Metazoan genes and overexpress post-Metazoan genes, underscoring the known alignment of ontogenesis with phylogenesis.

**Figure 7.**
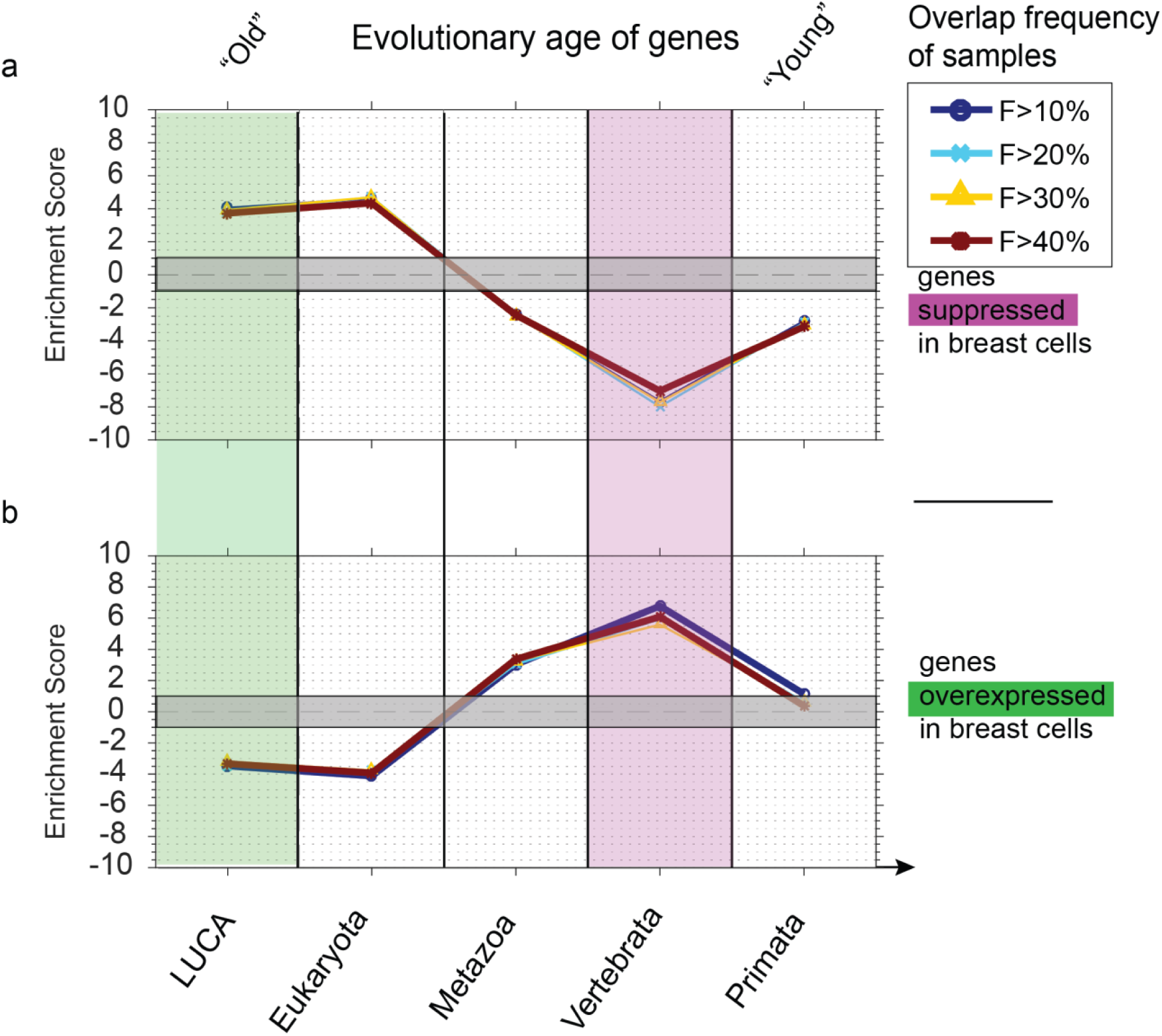
Evolutionary age enrichment analysis of genes suppressed and overexpressed in differentiated ductal breast cells relative to those in embryonic stem cells. (a) The frequency of the genes suppressed in breast cells were significantly enriched in LUCA and Eukaryota age groups and depleted in post-metazoan age groups. (**b**) The genes overexpressed in breast cells were significantly enriched in Metazoa and Vertebrata age groups and depleted in pre-metazoan age groups. The various overlap frequency categories F >10%, >20%, >30%, >40% did not affect the enrichment pattern due to the small variety of the cell samples.

## DISCUSSION

In recent years the classic somatic mutation theory has attracted much criticism due to the lack of evidence supporting it for large patient population data, such as the cancer genome project (TCGA). Accumulating evidence shows that non-genetic transitions of cell phenotypes and tumor micro-environment interactions play critical roles in tumorigenesis and cancer progression. The cancer atavism hypothesis proposed that cancer is a throwback to a (modified) primordial phenotypic state characterized by autonomous cell proliferation and increased resilience. By associating the genes differentially expressed between cancer tissues and their normal tissue counterparts with the genes’ evolutionary ages, we found that cancer cells tend to suppress post-Metazoan genes and overexpress pre-Metazoan genes – supporting the idea that cancer cells enter an ancient cell state which afford cells with enhanced functions of cell proliferation and survival while weakening cell differentiation and inter-cellular signaling functions.

Although a vast amount of cancer genomic information is currently available, deciphering such data to account for the complexities and intricacies of multicellular life has proven to be a difficult task. Indeed, the compartmentalization of genes into distinct biological functions, such as cell division, angiogenesis and invasions etc., could lead to the discovery of robust therapeutic targets. It is well appreciated that individual genes rarely convey the capacity to drive and coordinate the complex cellular functions required for cancer progression but that simultaneous, well-orchestrated actions of multiple genes is required. Even though the complexity could be overwhelming, to an extent as almost to defy the imagination that they evolved in Darwinian fashion by random mutation (accelerated by the cancer’s genome instability) and selection, we must remember that as “damaged” as they may be, cancer cells are still functional – unfortunately too robustly so. They recapitulate life forms that ought to behave and respond “normally” in many biologically relevant ways. Therefore, only a fraction of their functionality can possibly be impaired, these being mainly programs of communication and “decorum” in a collective context rather than core functions that determine the survival of individual cells. In particular, tumor samples of even the most similar histopathological subtypes generally present very different patterns of genetic alterations, yet the molecular pathways affected are typically quite similar^72^.

Our central idea is that all the relevant cellular alterations that drive cancer must disrupt the balance normally enforced by collective processes in the context of multicellular stability and homeostasis. In turn, such breakdown removes the collective regulations that would normally remove the abnormal cells affording them with newfound survival and replicative potential outside the multicellular ideology. This observation considerably limits the scope of biological processes that need to be considered. If we take into account the phenotype of multicellularity, two classes of general cellular processes should be playing out in balance: cell-level programs that determine the life cycle, replication and survival of individual cells, and collective-level programs that act at the organism level, determining its life cycle, reproduction and survival. The first kind of program provides regulations over cells; the second class provides regulations at the organismal level, but is still precluded by cell-level regulations. In this context, which alterations are “tolerated” by regulation controls and basic cellular survival principles would depend entirely on whether both cellular and organismal levels of regulations are or just cellular one in play.

Generally the processes involved in cancer progression are embroiled in the disruption of the equilibrium between these two forces. Here we extended this basic principle further to address gene sets in the context of molecular pathways. We distinguished between cell-promoting pathways, generally associated with rapid cell replication, early development, and DNA repair and survival response to environmental challenges on the one hand, and tissue-promoting pathways associated with cell-cell communication, suppression of cell replication by collective controls, morphogenesis and homeostasis on the other hand. Given that these two types of programs convey differential regulations to either cells or multicellular organisms, it can be expected that the former corresponds to ancient functions, relatively well conserved, while the latter corresponds to more modern functions associated with multicellularity. Our age enrichment and pathway analyses are consistent with and complement previous research associated with the cancer atavism hypothesis^38,41,43,45,46^. Furthermore, we looked at the microarray data of normal human breast cell differentiation and found the evidence that cell differentiation generally up-regulates post-Metazoan genes and down-regulates pre-Metazoan genes. Once gene expression profiles of various transient and terminal cell types during embryogenesis and development become available^73^, a systematic study of the gene evolutionary ages of all cell types, relative to embryo stem cells, will reveal if one can observe similar phenomena to that reported here and allowing us to map the entire cell developmental tree to phylogenesis in a systematic manner.

## Material and Methods

### Phylostratigraphy of human genes

The evolution ages of human genes were extracted from the eggNOG 4.0 database^58^. This database has over 7.7 million proteins categorized into 1.7 million orthologous gene families derived by functional annotations and genetic sequence overlaps, comprising more than 3,600 species. Initially, we identified the orthologous groups that contain human genes and downloaded the pre-computed phylogenetic trees. If the orthologous group included a single human gene, then the phylostratigraphic age was the most ancient phyla of species represented. If there are multiple human species, the phylogenetic trees where parsed at the last common ancestor node for each individual human genes. Each sub-tree is then used to assign the phylostratigraphic age.

The gene families provided by eggNOG are covered in eleven main taxonomic levels for the human lineage: Last Universal Common Ancestor (LUCA), Eukaryota, Opisthokonta, Metazoa, Bilateria, Chordata, Vertebrata, Mammalia, Euarchontoglires, Primata, and Hominidae. In order to have a good number of species per level (and have better statistical significance) we reduce these 11 levels into five basic levels (see the mapping table in Table 4):
1. Last Universal Common Ancestor (LUCA)
2. Eukaryota
3. MetazoaVertebrata
4. Primata

**Table 4.** Mapping table between 11-level evolution ages to 5-level evolution ages.

|  | **11-Level Age Names** | **5-Level Age Names** |
| --- | --- | --- |
| 1 | LUCA | LUCA |
| 2 | Eukaryota | Eukaryota |
| 3 | Opisthokonta | Eukaryota |
| 4 | Metazoa | Metazoa |
| 5 | Bilateria | Metazoa |
| 6 | Chordata | Metazoa |
| 7 | Vertebrata | Vertebrata |
| 8 | Mammalia | Vertebrata |
| 9 | Euarchontoglires | Vertebrata |
| 10 | Primata | Primata |
| 11 | Hominidae | Primata |

Genes from intermediate levels were included at a correspondingly lower accepted level. As a result of this methodology, we obtain the taxonomic level for each human gene in the database (19,177 genes). We refer to these levels in order to indicate the rank of ancestry of genes, pathways and functions involved in cancer, motivated by the observation that orthologous genes seem more likely to retain ancestral gene functions^74^.

### Cancer oncogene and tumor suppressor gene lists

We studied 1,895 genes whose mutations are causally implicated in cancer. These genes are annotated as either oncogenes or tumor suppressor genes accordingly if the evidence indicates that the mutation promotes or suppresses cancer cell growth. This list was compiled by merging the well documented Catalogue Of Somatic Mutations In Cancer (COSMIC)^56^, the more recent list of cancer driver genes published by Vogelstein’s team^72^ and the lists of tumor suppressor genes^75^ and oncogenes^57^ published by M. Zhao and collaborators. From these gene lists, only those genes that have associated EGGNog phylostratigraphic ages were chosen to yield 980 oncogenes and 915 tumor suppressor genes.

### Ten solid tumor RNA-Seq transcriptomes from TCGA

The gene expression data of RNA-Seq in RPKM (Reads Per Kilobase per Million mapped reads) were retrieved from The Caner Genome Atlas (TCGA) project data portal (https://gdc.nci.nih.gov/). We focused our analysis on ten cancer types: breast invasive carcinoma (BRCA), Uterine Corpus Endometrial Carcinoma (UCEC), Colon adenocarcinoma (COAD), Head and Neck squamous cell carcinoma (HNSC), Kidney renal clear cell carcinoma (KIRC), Kidney renal papillary cell carcinoma (KIRP), Liver hepatocellular carcinoma (LIHC), Lung adenocarcinoma (LUAD), Lung squamous cell carcinoma (LUSC), Thyroid carcinoma (THCA).

### Human embryonic stem cells, iPSCs and breast epithelial cells microarray transcriptomes

We curated microarray gene expression profiles from 50 samples of embryonic stem cells (ESC) from GEO (http://www.ncbi.nlm.nih.gov/geo/). Their GSE accession numbers are GSE6561, GSE7234, GSE7896, GSE9086, GSE9440, GSE9709, GSE9832, GSE9865, GSE9940, GSE12390, GSE13828, GSE14711, GSE14897, GSE15148, GSE16654 and GSE18679. We curated microarray gene expression profiles from 73 samples of iPS cells from GEO. Their GSE accession numbers are GSE9832, GSE9709, GSE9865, GSE12390, GSE12583, GSE13828, GSE14711, GSE15148, GSE16654 and GSE18111. We also curated microarray gene expression profiles from 159 samples of breast epithelial cell lines of primary cell cultures from GEO. Their GSE accession numbers are GSE10780, GSE9649, GSE12917 and GSE13671.

### The cancer-related pathways

The cancer-related pathway data was obtained from the KEGG PATHWAY Database^63^ (http://www.kegg.jp/). We downloaded human gene lists of the pathways associated with cancer (all pathways included under the KEGG category ID 05200 Pathways in Cancer) and a list of 14 essential housekeeping pathways. These two lists are not mutually exclusive. The list of pathways is shown in Table 1 and 2. Thus all genes involved in each pathway that have a corresponding EGGNog age are considered in our analysis.

### The method used to calculate gene age enrichment score

We define gene age enrichment score for an observed gene list as the adjusted residual^76^ calculated in the following way:
1. Calculate the distribution of ages of the background gene list, as the frequency *F*_bg_ (a) of genes in each age group *a*
2. Calculate the expected frequency of observed ages according to the usual null hypothesis in a contingency table:

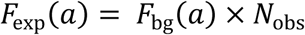

where N_obs_ is the number of genes in the observed list (e.g. tumor suppressor genes, oncogenes, etc.)
3. For each set 1,000 uniformly random samples of the same size as the observed list are drawn without replacement from the background list. Age frequencies 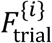 (*a*) are calculated for each trial *i*.
4. We use the distribution of frequency values for each age group across sample trials *G*_*a*_ *(F)* to statistically test the likelihood of observed frequencies.
5. In particular, it’s true that the expected value of the frequency for each age group *F*_*exp*_*(*a*;) = 〈 G*_*a*_*(F*_*i*_) 〉, with 〈*X*〉 the mean across trial samples.
6. The adjusted residual is the deviation of observed from expected frequencies normalized by 2 times the standard error of the bootstrapped trial samples *G*_*a*_ *(F*_*i*_) in each age group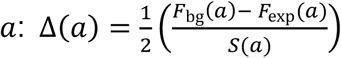 with 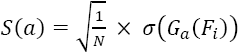 the standard error across trial samples and N = 1000 the number of trials.

This standardized measure takes into account the individual sizes of the samples^76^. The value of the adjusted residual can be directly interpreted in a probabilistic way, a value of ±1 corresponds to the standard 2-sigma test with p-value ~ 0.05. Hence adjusted residuals larger than 1 indicate that the observed value is larger than expected with statistical significance, i.e. the corresponding age group is over-represented, and equivalently adjusted residuals smaller than -1 indicate that the corresponding age group is under-represented. Thus the adjusted residual is used as a statistical score for gene age enrichment.

#### Acknowledgements

We thank Kimberly Bussey for helpful comments and discussions. We thank Merja Heinäniemi for helping the curation of gene expression profiles. This work was supported by National Institute of General Medical Sciences (NIGMS) Grant R01GM109964, NIGMS National Centers for Systems Biology Grant 2P50GM076547-06A1 and NIH Grant U54CA143682. This research was also supported by the National Science Foundation Grant PHY11-25915. The content is solely the responsibility of the authors. The funders had no role in study design, data collection and analysis, decision to publish, or preparation of the manuscript.

## Author Contributions

J.X.Z. and L.C. conceived the idea for the theoretical analysis, S.H. and P.D. conceived of the biological framework. J.X.Z., L.C., T.K. and K.T. curated data and performed data analysis. J.X.Z. and L.C. drafted the manuscript; P.D., I.S. and S.H. edited and wrote the paper.

## Data availability statement

The transcriptome data of 10 cancer types from TCGA are publicly available (https://portal.gdc.cancer.gov). The access IDs for different cancer types are: breast invasive carcinoma (BRCA), Uterine Corpus Endometrial Carcinoma (UCEC), Colon adenocarcinoma (COAD), Head and Neck squamous cell carcinoma (HNSC), Kidney renal clear cell carcinoma (KIRC), Kidney renal papillary cell carcinoma (KIRP), Liver hepatocellular carcinoma (LIHC), Lung adenocarcinoma (LUAD), Lung squamous cell carcinoma (LUSC), Thyroid carcinoma (THCA). The microarray data of Human embryo stem cells and ductal breast cells are publicly available. The access numbers are GSE6561, GSE7234, GSE7896, GSE9086, GSE9440, GSE9709, GSE9832, GSE9865, GSE9940, GSE12390, GSE13828, GSE14711, GSE14897, GSE15148, GSE16654, GSE18679, GSE12583, GSE18111, GSE10780, GSE9649, GSE12917 and GSE13671.

